# NSD1 supports cell growth and regulates autophagy in HPV-negative head and neck squamous cell carcinoma

**DOI:** 10.1101/2023.09.19.558537

**Authors:** Iuliia Topchu, Igor Bychkov, Demirkan Gursel, Petr Makhov, Yanis Boumber

**Affiliations:** Robert H. Lurie Comprehensive Cancer Center, Feinberg School of Medicine, Division of Hematology/Oncology, Northwestern University, Chicago, IL, 60611; Pathology Core Facility, Feinberg School of Medicine, Northwestern University, Chicago, Illinois, Chicago, IL, 60611; Cancer Signaling and Microenvironment Program, Fox Chase Cancer Center, Philadelphia, PA, 19111; Division of Hematology/Oncology, Sections of Thoracic / Head and Neck Medical Oncology, O’Neal Comprehensive Cancer Center, Heersink School of Medicine, University of Alabama in Birmingham, Birmingham, AL, 35233

**Keywords:** NSD1, Head and neck cancer squamous cell carcinoma (HNSCC), Akt/mTORC1, autophagy

## Abstract

Head and neck squamous cell carcinoma (HNSCC) is the sixth most common cancer worldwide. Despite advances in therapeutic management and immunotherapy, the five-year survival rate for head and neck cancer remains at ∼66% of all diagnosed cases. A better definition of drivers of HPV-negative HNSCC that are targetable points of tumor vulnerability could lead to significant clinical advances. NSD1 is a histone methyltransferase which catalyzes histone H3 lysine 36 di-methylation (H3K36me^2^); mutations inactivating NSD1 have been linked to improved outcomes in HNSCC. In this study, we show that NSD1 induces H3K36me^2^ levels in HNSCC, and that the depletion of NSD1 reduces HNSCC of cell growth *in vitro* and *in vivo*. We also find that NSD1 strongly promotes activation of the Akt/mTORC1 signaling pathway. NSD1 depletion in HNSCC induces an autophagic gene program activation, causes accumulation of the p62 and LC3B-II proteins, and decreases the autophagic signaling protein ULK1 at both protein and mRNA levels. Reflecting these signaling defects, knockdown of NSD1 disrupts autophagic flux in HNSCC cells. Taken together, these data identify positive regulation of Akt/mTORC1 signaling and autophagy as novel NSD1 functions in HNSCC, suggesting that NSD1 may be of value as a therapeutic target in this cancer.

## Introduction

Head and neck cancer squamous cell carcinomas (HNSCC) arise from the mucosal epithelium and develop predominantly in the oral cavity, pharynx, and larynx (1). HNSCC is the sixth most common cancer worldwide with about 900,000 new cases annually (2), including more than 60,000 in the United States (3). In the US, approximately 4% of the US population will be diagnosed with HNSCC, and 15,000 die from this disease each year, representing a significant healthcare problem(4). Despite advances in therapeutic management and immunotherapy, the five-year survival rate for head and neck cancer remains at 59-76% of all diagnosed cases; for disease diagnosed at an advanced stage, 5-year survival is 34-49% depending on types of head and neck cancer (4). Factors such as smoking and alcohol consumption are strongly associated with risk of HNSCC, while a growing number of cases are caused by infection with human papillomaviruses (HPVs) (5). HPV-positive HNSCC patients have a more favorable prognosis than HPV-negative patients (1), while HPV-negative HNSCC is associated with much worse outcomes (6). Better definition of the targetable oncogenic mechanisms of HPV-negative HNSCC could lead to significant clinical advances.

Deregulation of histone methylation, including lysine 36 dimethylation (K36me^2^) of histone H3, is sometimes detected during tumor development and progression (7). NSD1, NSD2, and NSD3 are key histone methyltransferases (HMTs) that catalyze mono- and dimethylation of H3K36 (8). Depletion of NSD1 is associated with DNA hypomethylation in HNSCC cells (9, 10), and NSD1 depletion usually leads to a significant reduction of level of H3K36me^2^ at the intergenic genomic regions (11). *NSD1* inactivating mutations have been reported to occur at a frequency at 10-13% and correlate with a better prognosis for HNSCC patients (9, 12, 13), especially among patients with HPV-negative laryngeal tumors (9), in analysis of a cohort of patients in which treatment with chemoradiation was common.

Besides these global chromatin effects related to HMT activity, it has been demonstrated in other cancer types that NSD1 regulates additional oncogenic signaling pathways, such as Nuclear Factor-kappa B (NF-κB) (14), Wnt/β-catenin (15, 16), and HIF1α (17). NSD1 loss decreases the growth of liver, breast and esophageal cancer cells (15–17), and NSD1 depletion leads to a moderate increase in sensitivity to cisplatin and carboplatin drugs (12, 13). Immune checkpoint inhibitors are showing great promise in numerous tumor types; however, HNSCC tumors with inactivating *NSD1* mutations have an “immune cold” microenvironment, characterized by the reduced infiltration by tumor-associated leukocytes (18). Further, a recent publication has used multiple *in vivo* and *in vitro* models to show that depletion of NSD1 leads to tumor immune evasion in HNSCC (19). Based on these results, it has been suggested that *NSD1*-mutated tumors may be less sensitive to immunotherapy, although data directly supporting this idea and details of the underlying mechanism are not yet available (18, 20). Overall, the mechanism by which NSD1 regulates cancer cell growth and therapeutic response is complex, and not fully understood.

The serine/threonine kinase mammalian target of rapamycin (mTOR) is a key regulator of cellular metabolism and a key driver of proliferation for many types of cancer (21). Activation of the PI3K-Akt-mTOR pathway occurs in the majority of HNSCC cases and contributes to tumor growth (22–24). The mTOR kinase is a core constituent of two protein complexes mTORC1 and mTORC2. mTORC1 promotes cell growth through the activation of the translation regulator S6 kinase (S6K), which in turn promotes protein synthesis (21). mTORC1 also inhibits autophagy (25), a process of recycling cell components that is usually considered as a protective mechanism for cancer cells growing in nutrient poor conditions that induce metabolic stress (26). mTOR inhibition through small molecule drugs typically initiates autophagy (25).

In this study, we have found that the striking reduction of HNSCC growth *in vitro* and *in vivo* induced by NSD1 depletion is accompanied by repression of the Akt/mTORC1 pathway. While this would normally induce autophagy, NSD1 knockdown blocks of the initial stages of autophagy through suppression of the expression of ULK1 protein, and disrupts autophagic flux, as reflected by although p62 and LC3B accumulation. These and other data suggest a dual role of the NSD1 protein in autophagy regulation: inhibition through the support of the mTORC1 pathway and activation of initial stages of autophagy via support of ULK1 expression. This work highlights a previously unsuspected role of NSD1 in control of cell growth and autophagy in cancer.

## Materials and Methods

### Cell lines and cell culture

Human HNSCC cell lines FaDu, Cal27, SSC61, SCC4 were obtained from the American Type Culture Collection (ATCC); JHU 011 and JHU 022 were gifts from Dr. E. Izumchenko (University of Chicago). A more detailed characterization of these cell lines is presented in Supplementary Table 1. HEK293T cells were used for retro- and lentivirus production. All cell lines were grown in RPMI-1640 with 10% FBS and 100 U/ml penicillin/streptomycin (Gibco). Cells were incubated in a 37 °C humidified incubator with 5% CO_2_.

### Vector construction and virus production

To generate stable cell lines with inducible NSD1 knockdowns, self-complementary single-stranded DNA oligos (Supplementary Table 2) were annealed and cloned into AgeI/EcoRI sites of Tet-pLKO-puro vector (Addgene, #21915). Tet-pLKO-puro vectors were packaged into a lentivirus system with pCMV-VSV-G (Addgene, #8454) and psPAX2 (Addgene, #12260). A pBABE-puro mCherry-EGFP-LC3B plasmid was obtained from Addgene (#22418) and packaged into a retrovirus system with packing plasmid pCMV-VSV-G (Addgene, #8454) and pUMVC (Addgene, #8449). HEK293T cell line was used for retroviral and lentiviral system amplification with TransIT-293 Transfection Reagent (Mirus).

### siRNA transfections

The sequences of siRNAs used for silencing NSD1 gene are shown in Supplementary Table 3; as a negative control we used siRNA Universal Negative Control #1 (SIC001, Sigma-Aldrich). Cells were plated onto 6-well plate for western blot or on glass coverslips for confocal microscopy. At 30% confluence cell were transfected with siRNAs at final concentrations of 20 nM using transfection agent TransIT-X2® Dynamic Delivery System (Mirus). Cells were lysed for western blot and/or fixed for confocal microscopy analysis and images acquisition, 72 hours post transfection.

### Total RNA-sequencing and Data Analysis

#### Total RNA-seq

mRNA was extracted from JHU 011 and Cal27 cell lines with NSD1 knockdown (pLKO was used as a control), post 72 hr of doxycycline induction. The stranded total RNA-seq was conducted at the Northwestern University NUSeq Core Facility. The Illumina Stranded Total RNA Library Preparation Kit was used to prepare sequencing libraries. The Kit procedure was performed without modifications. llumina HiSeq 4000 Sequencer was used to sequence the libraries with the production of single-end, 50 bp reads.

#### RNA-seq Analysis

The quality of reads, in FASTQ format, was evaluated using FastQC. Reads were trimmed to remove Illumina adapters from the 3’ ends using cutadapt (27). Trimmed reads were aligned to the *human* genome using STAR (28). Read counts for each gene were calculated using htseq-count (29) in conjunction with a gene annotation obtained from Ensembl (http://useast.ensembl.org/index.html). Normalization and differential expression were calculated using DESeq2 that employs the Wald test (30). The cutoff for determining significantly differentially expressed genes was an FDR-adjusted p-value less than 0.05 using the Benjamini-Hochberg method.

GSEA (Gene Set Enrichment Analysis) software (https://www.gsea-msigdb.org/gsea/index.jsp) was used for gene pathway analysis using gene sets from Molecular Signatures Database v7.5.1.

### RPPA Analysis

JHU 011 and Cal27 cell lines with NSD1 shRNA knockdown (pLKO is as a control, NSD1 sh1, NSD1 sh2) were lysed and prepared according to MD Anderson Core Facility instructions, as previously described, and RPPA was performed at the facility(31–33). Analysis was performed using 487 antibodies. Data were visualized using the GraphPad Prism software.

### Proliferation assay

500 cells/well were plated in 96-well cell culture plates in complete media. After 24h, NSD1 knockdown was induced by 1 μg/ml of doxycycline. CellTiter-Blue® assay (Promega) reagent was added and incubated for 1.5h, fluorescence was measured at 560/590 nm to obtain a 0h time point. Next, the procedure was repeated at 72, 96, 120, 148, and 168 hours. Proliferation was calculated as a relative value, where the 0h hour time point was taken as a one.

### Clonogenic assay

Cells were plated in 12-well plates (200 cells/well). After 24Uh, NSD1 shRNA knockdown or pLKO (control) were treated with 1 μg/ml of doxycycline to induce knockdown. Then cells were incubated for 7-10 days at 37 °C, 5% CO2 for colony formation. After 7-10 days cells were fixed with 10% acetic acid/10% methanol solution and stained with 0.5% (w/v) crystal violet. Plates were scanned and colonies were counted using Image J software.

### Apoptosis assay

Cells were seeded in a T-25 flask and NSD1 knockdown was induced for 144 hours. Next, cells were trypsinized, prepared according to the manufacturer’s protocol of the Alexa Fluor 488 Annexin V/Dead Cell Apoptosis Kit (Thermo Fisher Scientific), and analyzed using LSRFortessa Cell Analyzer (BD). At least 10U000 events were calculated for each sample.

### RNA isolation and RT-qPCR

For total RNA isolation, Quick-RNA Miniprep Kit (Zymo research) was used according to the manufacturer’s protocol. Complementary DNA was synthesized using iScript Reverse Transcription Supermix (Bio-Rad). Gene expression analysis was performed with QuantStudio 3 Real-Time PCR System (Applied Biosystem), using SYBR Green PCR master mix (Applied Biosystem); primer sequences are listed in Supplementary Table 4.

### Western blot

Cell pellets were prepared with lysis buffer (50mM Tris pH 7.6, 2% SDS) with Halt Protease & Phosphatase Inhibitor Cocktail (Thermo Scientific). Protein concentrations were measured using Pierce BCA assay (Thermo Scientific). Separated proteins were transferred on the PVDF membrane. The membranes were blocked with 1% nonfat milk in TBST for 1h at room temperature. After blocking, the membranes were incubated overnight at 4°C with primary antibodies listed in Supplementary table 5. Bands were developed using SuperSignal West Pico Plus Solution (Thermo Scientific) and detected with an autoradiography CL-Xposure Film (Thermo Scientific). Films were scanned and images were quantified with Image J software.

### Assessment of *in vivo* tumor growth

For in vivo tumor growth studies, 5□×□10^6^ of FaDu cells with inducible NSD1 shRNA knockdown (FaDu pLKO is a control) were injected subcutaneously in the flank region of 7-week-old mice using CB17-SCID mice (Charles River), with 8 mice per each group. All animal procedures were done using institutionally approved animal protocol. After 5 days of tumor growth, when tumors volume reached 30-40 mm^2^, mice have been started to feed with Doxycycline Rodent Diet (200 mg/kg, Bio-Serv) to induce shRNA knockdown. Tumors were measured every 5 days. Tumor volume was calculated with the formula: [volume□=□0.52□×□(width)^2^□×□length]. After 25 days of the post-tumor cells injection, mice were euthanized, and tumor tissues were collected for histology and western blot.

### Immunohistochemistry of mouse xenografts

Tumor tissues from mice were collected, fixed in 10% phosphate-buffered formaldehyde (formalin) for 36 hours, and submitted to Mouse Histology & Phenotyping Laboratory (MHPL) of Northwestern University. Samples were embedded in paraffin, then hematoxylin and eosin (H&E) staining, immunohistochemistry (IHC) were performed by using standard protocols. Antibodies for p62 and LC3B proteins detection and their dilutions used for IHC are listed in **Supplementary table 5**. Slides were scanned and H-score was calculated using QuPath software (https://qupath.github.io/).

### Tissue Microarray Construction and Immunohistochemistry

Head and Neck surgical specimens resected at Northwestern Memorial Hospital, including were used to construct tissue microarrays (TMA). Clinical information (**Supplementary Table 6**) was available from the repository database and abstracted from clinical databases in an anonymized fashion (Northwestern IRB project # STU00214658, approved on 05/03/2021). The map of TMAs was created, reviewed, and constructed with the size of 1.5 mm core by using the semi-automatic Veridiam Tissue Microarrayer VTA-100. Immunohistochemical studies were performed on 4-micron sections from Formalin-Fixed Paraffin Embedded (FFPE) TMA blocks on charged slides by using Leica Bond-Max Autostainer. Antibody dilutions used are listed in **Supplementary table 5.** Stained slides were scanned using digital slide scanner Nanozoomer 2.0-HT (Hamamatsu); H-score was calculated using QuPath software (https://qupath.github.io/).

### ChIP-qPCR

Approximately 3 × 10^7^ cells were cross-linked with 1% formaldehyde. Chromatin immunoprecipitation (ChIP) assay was performed by using SimpleChIP Enzymatic Chromatin IP Kit (Cell Signaling Technology, #9003) according to the manufacturer’s protocol. For immunoprecipitation, 2,5 μg of rabbit anti-H3K36me^2^ antibody (Abcam, ab9049) was used per IP; the same amount of normal rabbit IgG antibodies was used as a control (CST, #2729) Purified DNA from immunoprecipitated chromatin was subjected to qPCR analysis using sing SYBR Green PCR master mix (Applied Biosystem). Primer sequences for *ULK1* gene regions are listed in Supplementary Table 7. The results were calculated as a fold enrichment by normalizing on IgG control.

### Autophagic flux measurement

Autophagy degradative activity (autophagic flux) was measured using an expression vector encoding the fusion protein mCherry-EGFP-LC3B (Addgene, #22418). We transfected JHU 011 cell line with pBabe-mCherry-EGFP-LC3B vector and established a stable cell line by puromycin selection. Cells were plated on the glass cover sleeps and NSD1 was knock downed with siRNA for 72 hours. Then cells were fixed in 4% PFA for 20 minutes and mounted in ProLong Gold Antifade Reagent with DAPI (Invitrogen). Cells were visualized using a NIKON A1R (B) GaAsP confocal microscope fitted with a 100×, 1.4-NA objective in the presence of immersion oil. Images were analyzed in the Fuji software using Analyze Particles plugin to calculate mCherry^+^ and EGFP^+^ single puncta and mCherry^+^ EGFP^+^ double-positive puncta. Puncta were quantified as a percent ratio per cell (n = 28 cells per condition).

### Statistical methods

All used statistical analyses noted in the figure legends were performed and visualized using GraphPad Prism software (v.9.5.1)

## Results

### NSD1 regulates H3K36me^2^ level along with cell proliferation and tumor growth in HNSCC

First, we evaluated the NSD1 and H3K36me^2^ in human normal epithelial tissues and in primary HNSCC samples. Tissue microarrays (TMAs) using with HNSCC tumor tissues derived from 36 HPV-negative patients revealed that both NSD1 and H3K36me^2^ protein expression were significantly lower in normal epithelium tissues compared to HNSCC tumor tissues of stages II through IV (**Figure 1A and B**).

**Figure 1.**
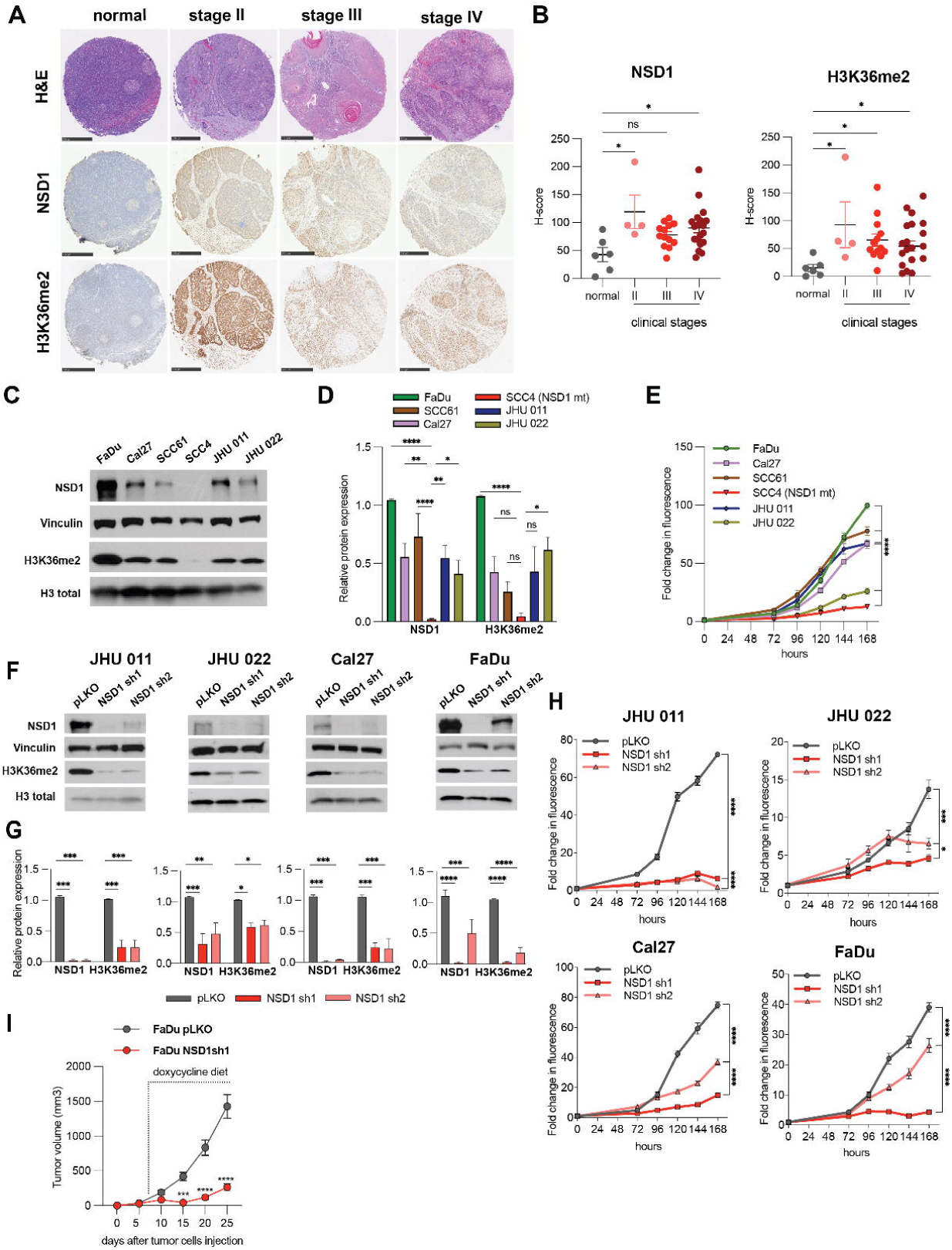
NSD1 regulates H3K36me^2^ level along with cell proliferation and tumor growth in HNSCC. (A) Representative images of human tumor tissues stained with NSD1 and H3K36me^2^ antibody. Scale bars, 500 μm. (B) H-score calculations from (A). Statistical significance was determined by Kruskal-Wallis with Dunn’s multiple comparisons post-test. (C) Western blot of NSD1 and H3K36me^2^ levels in a panel of human head and neck cancer cell lines. (D) Quantification of Western blot images in (C). Statistical significance was determined by ANOVA with Dunnett multiple comparison post-test. Each group was compared to SCC4 cell line. (E) Proliferation HNSCC cell lines as measured by CTB for 168 hours. Statistical significance determined by ANOVA with Dunnett multiple comparison post-test. Each group was compared to SCC4 cell line. (F) Western blot of NSD1 and H3K36me^2^ protein levels in NSD1 shRNA knockdown cells at 72h after knockdown induction. (G) Quantification of Western blot images in (F). Statistical significance was determined by ANOVA with Dunnett multiple comparison post-test. Each group was compared to pLKO. (H) Proliferation of pLKO-transfected or NSD1 shRNA transfected cell lines post doxycycline induction, as measured by CTB assay for up to 168 hours, at indicated time points. Statistical significance was determined by ANOVA with Dunnett multiple comparison post-test. Each group was compared to pLKO. (I) Quantitation of tumor volume in mice subcutaneously injected with FaDu NSD1 knockdown cell line (FaDu pLKO as a control); n=8 per group at indicated time points. Mouse number #1 from the control group died on day 23, tumor measurements were included in calculations until day 20. Statistical significance was determined by ANOVA with Šidák multiple comparison post-test. Experiments were performed in at least three independent biological repeats. The error bars are presented as mean ± SEM. ns – not significant, *p<0.05, **p<0.01, ***p<0.001, and ****p<0.0001.

Next, we examined the baseline expression levels of NSD1 and H3K36me^2^ in six HPV-negative HNSCC cell lines. The western blot quantification demonstrates that the highest level of NSD1 protein is in the FaDu cell line, while the SCC4 cell line carrying the frameshift loss of function mutation of NSD1 has the lowest level of this protein (**Supplementary table 1, Figure 1C and D**). mRNA expression level of NSD1 was also decreased in SCC4 cells compared to other HNSCC cells (**Figure S1A**). NSD1 has two structural and functional paralogs with partially redundant function: NSD2 and NSD3 (8). Therefore, we analyzed whether NSD2 and NSD3 mRNA and protein levels may correlate with NSD1 expression levels in these cell lines. As demonstrated in **Supplementary Figure S1B-D**, there was no such correlation indicating the fact that NSD1/2/3 proteins express independently of each other in HNSCC.

We then evaluated proliferation rate in six HNSCC cell lines. **Figure 1E** demonstrates that SCC4 had the lowest basal level of proliferation, while FaDu cells, with the highest NSD1 expression level, showed notably high proliferation rate. Taken together these data pointed out that the NSD1 expression level may have an impact on HNSCC cells growth.

Therefore, we investigated the role of NSD1 depletion in the isogenic cell line models. Thus, we developed four HPV-negative cell line models supporting doxycycline-inducible shRNA knockdown of NSD1 (in JHU 011, JHU 022, Cal27, and FaDu cells), each using two independent shRNAs against NSD1, with corresponding control models containing an empty pLKO as an empty vector. Interestingly, NSD1 depletion in those cells resulted in dramatic decrease of the H3K36me^2^ levels (**Figure 1F and G**). In parallel, we confirmed that neither the mRNA nor protein expression of NSD2 and NSD3 was affected in NSD1-depleted cells, excluding possible off-target effects of the shRNAs, or potential effects of NSD1 knockdown on the expression on its paralog proteins (**Supplementary Figure S1E-G**). We then evaluated the effect of NSD1 depletion on cell proliferation using CTB assay. Importantly, NSD1 knockdown resulted in potent suppression of cell growth in all four models (**Figure 1H**). Clonogenic analysis also demonstrated that NSD1 loss significantly reduced the ability to form colonies (**Supplementary Figure S2A**). Further, at six days after NSD1 knockdown, the level of apoptosis in the JHU 011 and Cal27 lines was slightly higher compared to control cells (**Supplementary Figure S2B**). This points out that NSD1 loss may promote apoptosis induction in HNSCC).

Next, to expand our *in vitro* observations supporting our hypothesis that NSD1 activity has an important role in regulation of HNSCC cells growth, we performed *in vivo* studies. To analyze the effects of NSD1 knockdown, FaDu cells bearing empty pLKO vector or expressing NSD1 shRNA (sh1) were subcutaneously injected into CB17-SCID mice, and NSD1 knockdown was induced by doxycycline 5 days after tumor cell injection. Excitingly, NSD1 depletion resulted a dramatic suppression of tumor growth (**Figure 1I**). We also confirmed efficient depletion of NSD1 depletion, and reduced H3K36me^2^ levels, in collected tumors (**Supplementary Figure S2C and D**).

Collectively, our data clearly demonstrate that NSD1 positively regulates H3K36me^2^ level in HNSCC and functionally supports HNSCC cells proliferation and tumor growth, both *in vitro* and *in vivo*. We were also able to demonstrate a significant up-regulation of the levels of H3K36me^2^ and NSD1 in HNSCC clinical samples compared to normal epithelial tissues.

### NSD1 depletion leads to accumulation of p62 and LC3B-II proteins in HNSCC, suggesting autophagy disruption

To explore which cell signaling programs are affected by NSD1 depletion, we performed RNA sequencing (RNA-seq) in JHU 011 and Cal27 cells, comparing cells 72 hours after induction of NSD1 shRNA or control. Gene set enrichment analysis (GSEA) indicated that NSD1 knockdown cells upregulated gene programs related to autophagy and response to starvation program **(Figure 2A and Supplementary Figure S3**).

**Figure 2.**
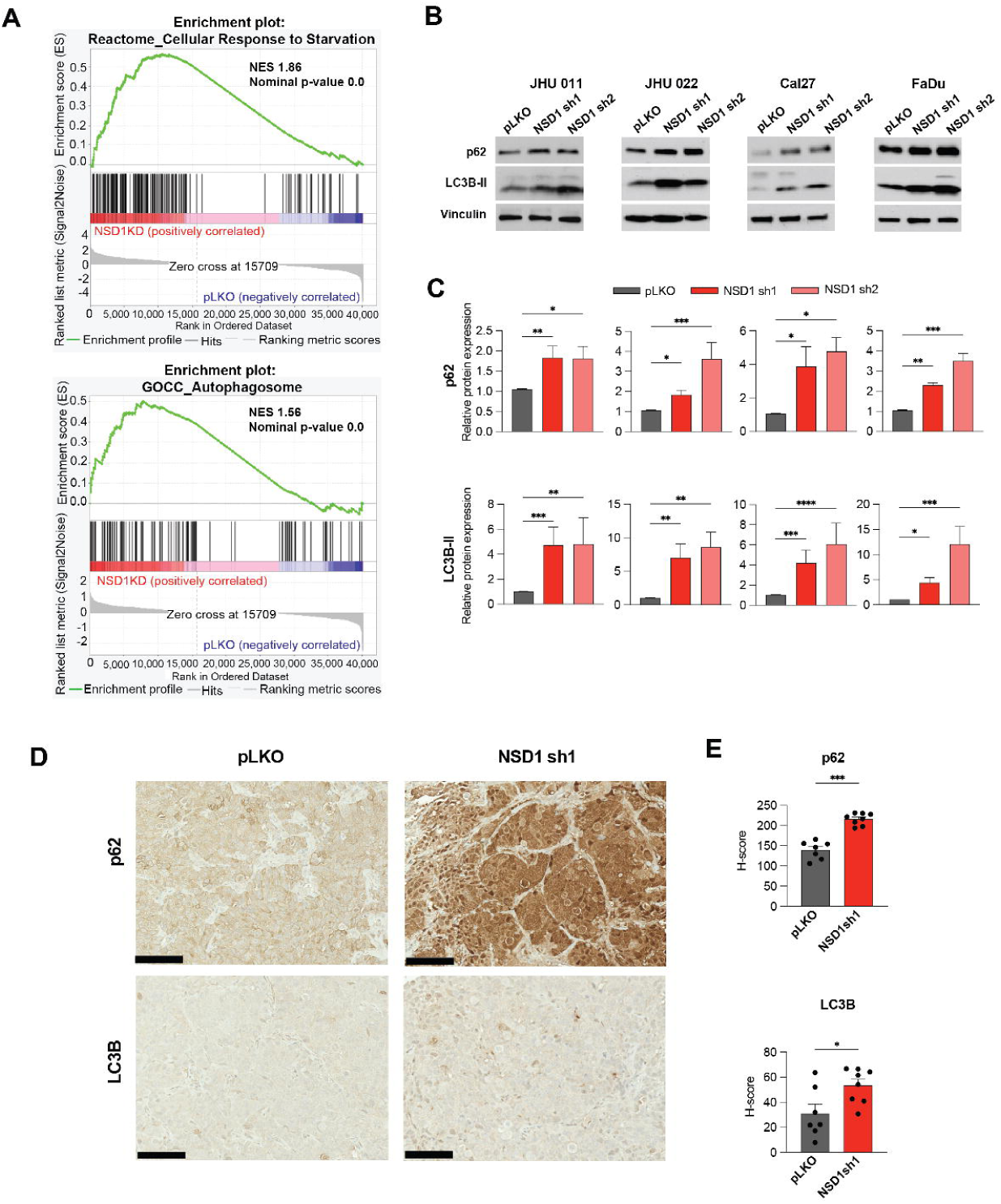
NSD1 depletion leads to accumulation of p62 and LC3B-II proteins in HNSCC, suggesting autophagy disruption. (A) Normalized enrichment score (NES) for genes identified as part of the Reactome_Cellular Response to Starvation and GOCC_Autophagosome gene sets after GSEA of RNA-seq in JHU 011 and Cal27 cell lines. (B) Western blot of p62 and LC3B-II protein levels in JHU 011, JHU 022, Cal27, and FaDu cell lines upon doxycycline induction of NSD1 shRNA knockdown or pLKO control. (C) Quantification of western blot images in (B). Statistical significance was determined by Kruskal-Wallis with Dunn’s multiple comparisons post-test. (D) Representative images for p62 and LC3B IHC staining in FaDu pLKO and NSD1sh1 tumors from the mouse xenograft models (20x magnification, scale bar 100 μm). (E) Quantification of average H-score from (D). Statistical significance was determined by Mann-Whitney test. Experiments were performed in at least three independent biological repeats. The error bars are presented as mean ± SEM. ns – not significant, *p<0.05, **p<0.01, ***p<0.001, and ****p<0.0001.

To confirm that NSD1 depletion affects autophagy, we evaluated LC3B-II and p62 protein levels by western blot in HNSCC cell lines. Expectedly, all tested cell lines demonstrated a significant accumulation of LC3B-II and p62 proteins after 72 hours of NSD1-knockdown (**Figure 2B and C**). The IHC staining of subcutaneous FaDu tumor tissues (**Figure 1I**) demonstrated potent accumulation of p62 and LC3B-II (**Figure 2D and E**). Taken together, our findings suggest disruption of autophagy upon NSD1 depletion in HNSCC.

### Depletion of NSD1 inhibits Akt/mTORC1 signaling pathway

Next, to explore the effects of NSD1 loss on HNSCC cell signaling pathways, we performed reverse phase protein array (RPPA) analysis in JHU 011 and Cal27 cell lines 72 hours after induction of NSD1 shRNA or control (pLKO). The RPPA data clearly demonstrated a decrease in activity of Akt/mTORC1 signaling pathway in NSD1 depleted HNSCC cells (**Figure 3A and B**).

**Figure 3.**
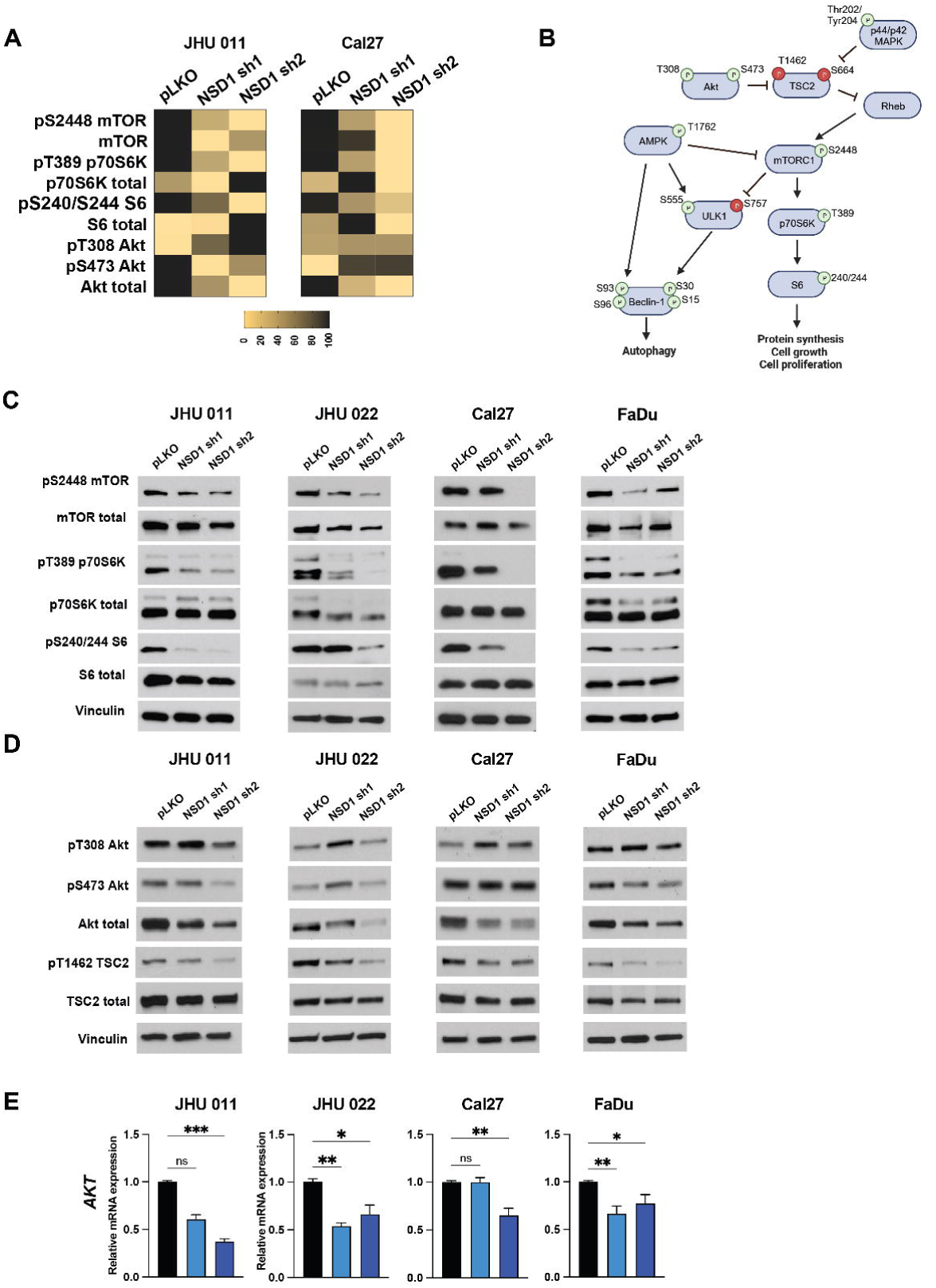
Depletion of NSD1 inhibits Akt/mTORC1 signaling pathway. (A) A Heatmap of the RPPA results for JHU 011 and Cal27 cell lines. (B) Schema of Akt/mTORC1 signaling pathway. (C) Western blot of mTOR and its downstream target protein levels in HNSCC cell lines with induced NSD1 shRNA knockdown. (D) Western blot of Akt and its TSC2 protein levels in HNSCC cell lines with induced NSD1 shRNA knockdown. (E) mRNA expression of the *AKT* gene in HNSCC cell lines post doxycycline induction of pLKO control or NSD1 shRNA knockdown, as measured by qRT-PCR. *Akt* relative level was normalized to *18S*, a control gene. Statistical significance was determined by Kruskal-Wallis with Dunn’s multiple comparisons post-test. Experiments were performed in at least three independent biological repeats. The error bars are presented as mean ± SEM. ns – not significant, *p<0.05, **p<0.01, ***p<0.001, and ****p<0.0001.

Further validation of the RPPA data has confirmed our observations. Indeed, NSD1 knockdown reduces activity of mTORC1 downstream signaling, as documented by decreased levels of phosphorylation levels of mTOR, p70S6K and S6 proteins in all tested HNSCC cell lines (**Figure 3C and Supplementary Figure S4**). Given that Akt is an upstream activator of mTORC1, which stimulates mTORC1 through inhibitory phosphorylation of TSC2 (**Figure 3B**) (21), we evaluated the effects of NSD1 depletion on Akt and its downstream effectors. As demonstrated in the **Figure 3D, E and Supplementary Figure S5A,** we observed a dramatic decrease of both, protein and mRNA levels of Akt. Importantly, this was accompanied with the reduction of TSC2 phosphorylation (T1462). Another well-known upstream effector of TSC2 protein, p44/42 MAPK, also demonstrated significant changes in total isoform level in JHU 011, Cal27, FaDu, but not in JHU 022 cells (**Supplementary Figure S5B-C).**

### NSD1 affects the initial stages of autophagy through the direct regulation of *ULK1* gene expression in HNSCC

Given that mTORC1 activity negatively regulates autophagy activation, we expected a mechanistic connection between downregulation of mTORC1 signaling and activation of autophagy-related gene programs upon NSD1 depletion.

Surprisingly, we observed that ULK1 total protein and mRNA expression levels were reduced (**Figure 4A-C**) in all tested HNSCC cell lines. This suggests the direct NSD1 control of *ULK1* gene expression via H3K36me^2^-dependent mechanism, as it was shown for other NSD1-regulated genes (19, 34, 35). To understand molecular mechanism of ULK1 regulation, we performed ChIP-qPCR analysis. NSD1-depleted cells demonstrated dramatic reduction of H3K36me^2^ level in the region of *ULK1* promoter, suggesting direct regulation of *ULK1* gene expression by NSD1 (**Figure 4D and E**).

**Figure 4.**
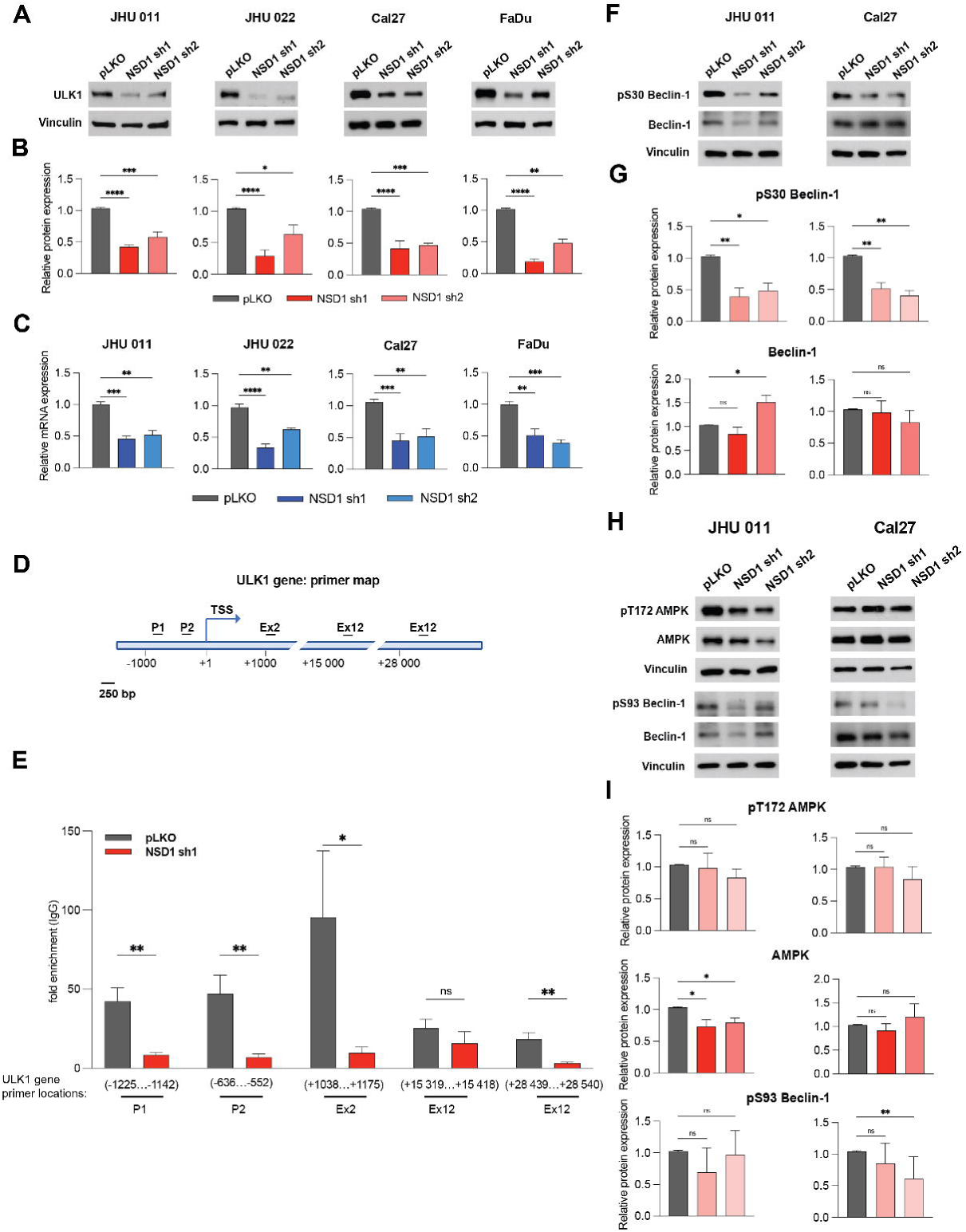
NSD1 affects the initial stages of autophagy through the direct regulation of *ULK1* gene expression in HNSCC. (A) Western blot of ULK1 protein level in JHU 011, JHU 022, Cal27, and FaDu cell lines upon doxycycline induction of NSD1 shRNA knockdown or pLKO control. (B) Quantification of Western blot in (A). Statistical significance was determined by Kruskal-Wallis with Dunn’s multiple comparisons post-test. (C) mRNA expression of the *ULK1* gene HNSCC cell lines post doxycycline induction of pLKO control or NSD1 shRNA knockdown, as measured by qRT-PCR. *ULK1* relative level was normalized to *18S* as a control gene. Statistical significance determined by Kruskal-Wallis with Dunn’s multiple comparisons post-test. (D) Primer location map of the *ULK1* gene regions for CHIP-qPCR. P – promoter, ex – exon, TSS – transcription start site. (E) CHIP-qPCR with H3K36me2 antibodies on the *ULK1* gene. Statistical significance was determined by multiple Mann-Whitney test. (F) Western blot of pS30 Beclin-1 and Beclin-1 total protein levels in JHU 011 and Cal27 cell lines upon NSD1 knockdown upon doxycycline induction of NSD1 shRNA knockdown or pLKO control. (G) Quantification of Western blot in (D). Statistical significance was determined by Kruskal-Wallis with Dunn’s multiple comparisons post-test. (H) Western blot of pT172 AMPK, AMPK total, pS93 Beclin-1, and Beclin-1 total protein levels in JHU 011 and Cal27 cell lines upon NSD1 knockdown upon doxycycline induction of NSD1 shRNA knockdown or pLKO control. (I) Quantification of Western blot in (H). Statistical significance was determined by Kruskal-Wallis with Dunn’s multiple comparisons post-test. Experiments were performed in at least three independent biological repeats. The error bars are presented as mean ± SEM. ns – not significant, *p<0.05, **p<0.01, ***p<0.001, and ****p<0.0001.

Moreover, the phosphorylation of Beclin-1 (pS30), downstream target of ULK1 (36), was decreased in NSD1-depleted cells of JHU 011 and Cal27 cell lines. The level of total isoform of Beclin-1 was not changed significantly, demonstrating trend to increase only under one of the shRNA in JHU 011 cell line (**Figure 4F and G**). Given that AMPK can activate Beclin-1 by direct phosphorylating at Serine 93 (37, 38), we evaluated the potential autophagy-activating compensatory effect of AMPK upon the decrease of pS30 Beclin-1 phosphorylation caused by the downregulation of UKL1 expression. We observed a minor decrease of AMPK protein level accompanied with the insignificant change in pS93 Beclin-1 phosphorylation, which only trended to decrease in JHU 011 cell line. Cal27 cell line didn’t demonstrate any change in phosphorylated and total AMPK protein levels but had a slight decrease in pS93 Beclin-1 protein (**Figure 4H and I**). Therefore, it led us to the conclusion that the decrease in the ULK1 level has not been compensated by AMPK activity. Collectively, these results indicate that NSD1 positively directly regulates ULK1 gene expression, and that the depletion of NSD1 may promote inhibition of the autophagy cascade at the initial stages in HNSCC.

### Autophagic flux is disrupted in NSD1-depleted HNSCC cells

The above findings clearly demonstrated that NSD1 depletion in HNSCC cells promoted the accumulation of p62 and LC3B-II, but also resulted in the decrease of ULK1 levels and pS30 Beclin-1 phosphorylation, which might seem to be contradictory results.

To explore whether NSD1 knockdown leads to initiation of autophagy or a block in autophagic flux (autophagy degradative activity system), we transfected JHU 011 cell line wild type with pBabe-mCherry-EGFP-LC3B plasmid (**Figure 5A**) and performed NSD1 knockdown using two different siRNA (**Figure 5B and C**). We observed the increase in the percentage of yellow puncta (mCherry+EGFP+) in cells upon siNSD1 knockdown and simultaneously, the decrease in free red puncta (mCherry+EGFP-), which suggests a decrease in autophagy flux efficiency (**Figure 5D and E**). Taken together, these results suggest that NSD1 supports the autophagy degradation activity. Depletion of NSD1 has led to the accumulation of autophagosomes, which was observed by western blot (evidenced by accumulation of p62 and LC3B-II) and visualized by LC3B-tagged protein, suggesting disruption of autophagic flux.

**Figure 5.**
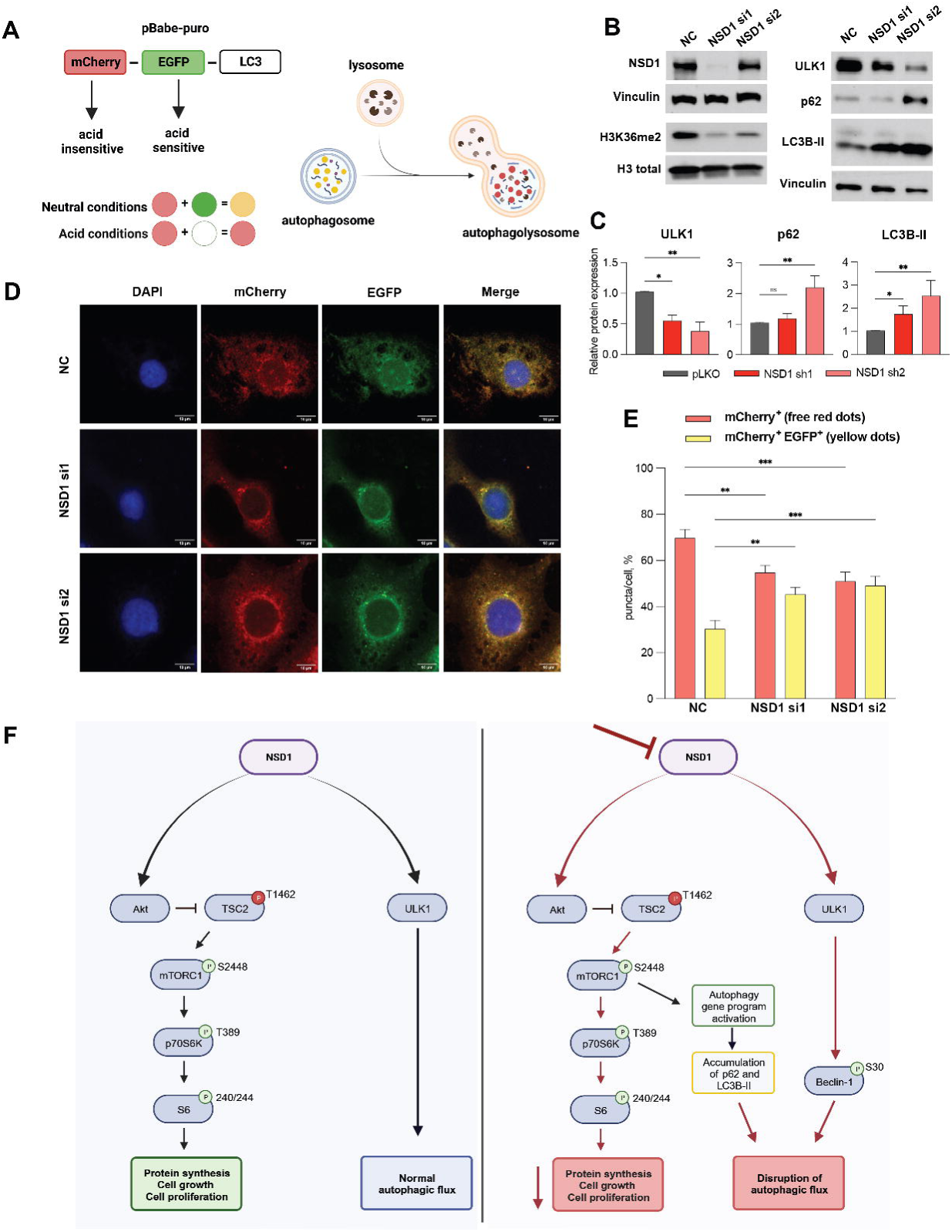
Autophagic flux is disrupted in NSD1-depleted HNSCC cells. (A) Schema of the mCherry-GFP-LC3 reporter to monitor autophagic flux: pBabe-mCherry-EGFP-LC3B reporter where the GFP tag is acid-sensitive while the mCherry tag is acid-insensitive. That means, that at the higher level of autophagic flux, autophagosomes fuse with the lysosome (which has acid pH) and the intensity of the EGFP tag will die out, while at the low level autophagic flux, both EGFP and mCherry tags will be detected and merged, resulting in yellow dots. (B) Western blot of NSD1, H3K36me^2^, ULK1, p62, and LC3B protein levels upon siRNA knockdown or negative control (NC) siRNA at 72hrs. (C) Quantification of western blot images in (B). Statistical significance was determined by Kruskal-Wallis with Dunn’s multiple comparisons post-test. (D) Representative confocal images of JHU 011 cells transfected with the pBabe-puro-mCherry-EGFP-LC3B, and with siRNA knockdown of NSD1 that was induced during 72 hours before cell fixation. Scale bars, 10 μm. (E) Quantification of puncta color percentage. Total number of autophagosomes (yellow) and autolysosomes (red) were quantified per cell (n=28 cells/condition). Statistical significance was determined by ANOVA with Dunnett multiple comparison post-test. Each group was compared to negative control (NC). (F) Schema of the proposed working model. Experiments were performed in at least three independent biological repeats. The error bars are presented as mean ± SEM. ns – not significant, *p<0.05, **p<0.01, ***p<0.001, and ****p<0.0001.

## Discussion

In the past decades, NSD1/2/3 histone methyltransferases have been shown to play an important role in hematologic malignancies and solid tumors (8). NSD1 is known to be oncogenic in leukemia, where NSD1 translocation drives a subset of acute myeloid leukemias with t(5;11)(q35;p15.5) which results in NUP98–NSD1 fusion protein (35). The role of NSD1 in solid tumors is less well-established. In particular, while a subset of HPV-negative HNSCC, primarily laryngeal carcinoma with *NSD1* loss of function mutations, has a favorable prognosis (9), this represents only a minority of HNSCC, while >80% HPV-negative head and neck tumors are *NSD1*-wild type. Interestingly, a recent Gameira et al paper showed that while in HPV+ HNSCC, NSD1/2/3 low expression cases associate with inferior outcomes, NSD1 and NSD2 high expressing HPV-negative HNSCC tumors show significant trends to associate with poor outcome (39). Therefore, in this study, we tested the hypothesis that elevated levels of NSD1 may be oncogenic and could promote tumor progression in HPV-negative HNSCC. We compared HNSCC cell growth in WT vs mutant cells lines, and *NSD1*-mutant cell line with the lowest NSD1 expression levels was the slowest growing, suggesting that NSD1 may sustain HNSCC cell growth. Next, we used NSD1-expressing HNSCC (laryngeal, tongue/hypopharynx, HPV-negative models) cell lines to test an impact of depletion of NSD1 on cell growth. HNSCC cell lines demonstrated dramatically reduced growth upon NSD1 depletion *in vitro*, and also *in vivo* using xenografts.

Our data contrasts somewhat with a recently published study by Li et al. which showed that the immunocompetent allograft HNSCC mouse model does not demonstrate the difference between WT and NSD1-knockout tumor growth; the authors explain this effect by the ability of NSD1 to affect the tumor-immune microenvironment (19). Additionally, it is certainly possible that mouse and human NSD1 roles in HNSCC could be different. Taken together, our findings in several independent HNSCC cell line models suggest that NSD1-expressing human HNSCC is dependent on NSD1 expression and suggests previously unsuspected oncogenic role for this enzyme in a large subset of HPV-negative HNSCC. Our findings complement advances in understanding the significance of H3K36 methyltransferases in HNSCC.

It was previously demonstrated that the NSD2 and NSD3, other members of NSD family proteins, have been implicated in HNSCC and their loss leads to the decrease in cell viability (40, 41), but here, after studying NSD1, we find that this enzyme is also critical for HNSCC cell growth. Farhangdoost et al recently demonstrated that CRISPR/Cas9 -generated knockout of *NSD1* as well as a mutation of *NSD1* in HNSCC cell lines may downregulate several gene programs including mTORC1 signaling (11). In our study, for the first time, we show the decrease of Akt/mTORC1 signaling in HNSCC, in response to NSD1 knockdown.

Here, we performed RNA-seq in JHU 011 and Cal27 with NSD1 shRNA knockdown cell lines allowed us further dissect mechanisms of HNSCC cell growth regulation by NSD1, and we found significant upregulation of starvation and autophagy-related gene expression programs. We suggested the autophagy is activated upon NSD1 knockdown in response to the mTORC1 pathway downregulation. Surprisingly, in the light of the fact that the activation of the gene program responsible for the for autophagosomes formation was upregulated after NSD1 knockdown, ULK1 gene was consistently positively regulated by NSD1 at both mRNA and protein levels. Some studies before described the epigenetic regulation of ULK1 by histone-methylation modifiers (42, 43). Here, we showed that mechanistically, ULK1 expression is directly and positively regulated by NSD1 and K36me^2^.

The increase in LC3B-II, along with accumulation of p62 protein in NSD1-depleted HNSCC cells indicates a failure to clear autophagosomes by fusion with the lysosomes. We confirmed autophagy defects by conducting autophagic flux experiments. Our observation of a significant decrease in the percentage of red fluorescence and increase in yellow fluorescence upon NSD1 depletion implies slowing down the autophagic flux. Taken together, decrease in Akt/mTORC1 activity and autophagy defects we observed suggests enhanced initiation of autophagy, which is ineffective in the absence of NSD1 (**Figure 5F**).

To our knowledge, NSD1/H3K36me^2^ control of autophagy has never been described before and is a novel and interesting finding. Inhibition of autophagy in cancer remains an attractive field, and several autophagy inhibitors are being tested in clinical trials (44). Similarly, PI3K/AKT and mTOR pathways are important for majority of HNSCC tumors because of several driver mechanisms, and clinical trials with agents targeting these pathways in HNSCC are ongoing (24, 45). In summary, we conclude that NSD1 enzyme is oncogenic and contributes to cancer cell growth in HPV-negative HNSCC, via support of Akt/mTORC1 pathway and regulation of autophagy. Inhibiting NSD1 enzyme and/or NSD1 downstream Akt/mTORC1 signaling, and to target/disrupt autophagy could be an exciting novel therapeutic strategy in HPV-negative HNSCC.

## Acknowledgments

We thank Dr. Erica Golemis for the constructive comments on the manuscript. We also thank Dr. Evgenii Izumchenko for the gift of JHU011 and JHU022 cell lines.

## Contributions

Conceptualization: IT, YB.; design and experiment performing: IT, IB, and DG; analysis of the data, visualization: IT and IB; original draft preparation: IT, PM, and YB. All Authors read and approved the final manuscript.

## Data availability

Data presented in this study are available from the corresponding authors upon request.

---

**Supplementary Figure S1.**
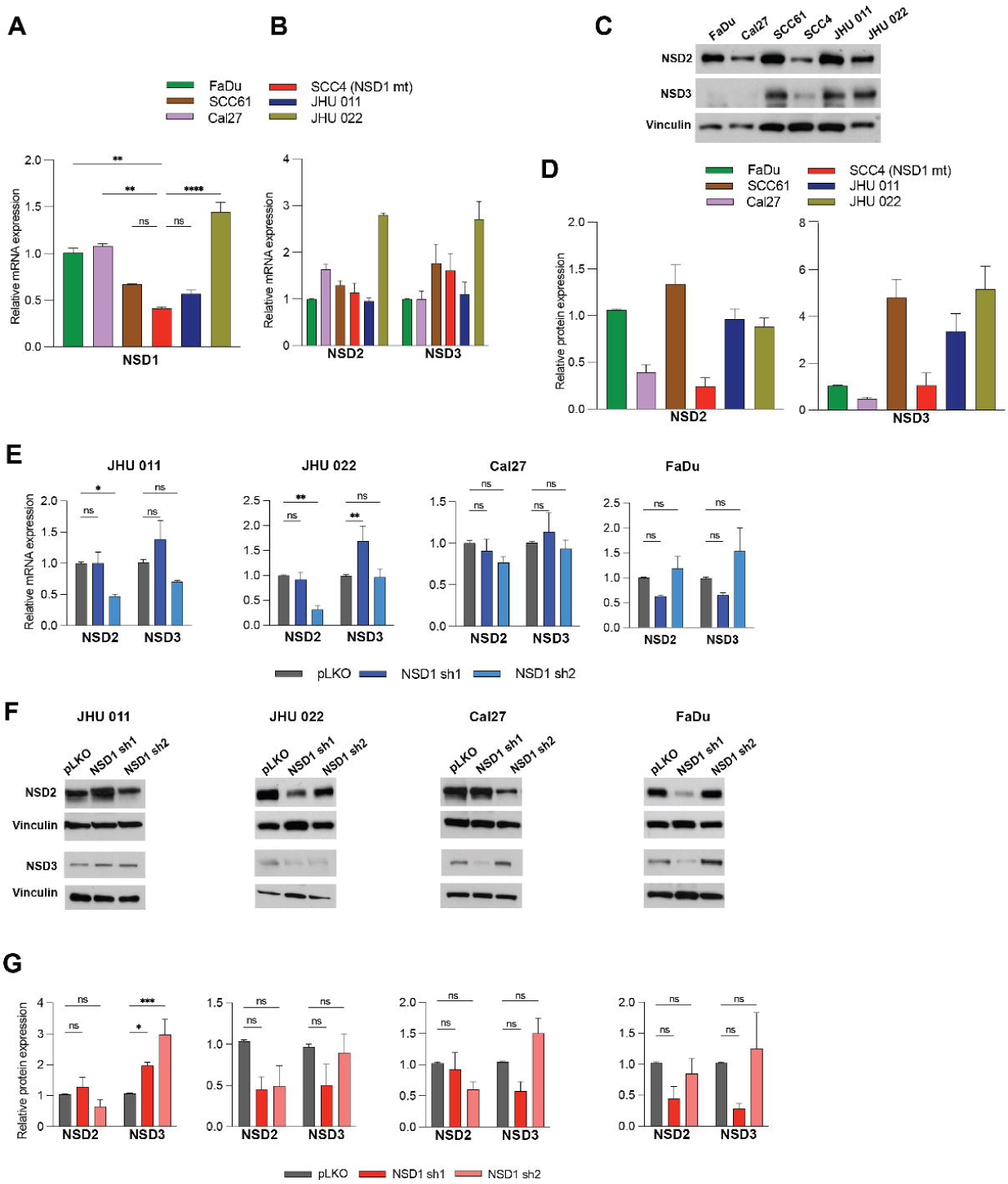
(A) Relative mRNA level of *NSD1* gene, measured by RT-qPCR in human HNSCC cell lines. *NSD1* relative level was normalized on *18S* as a control gene. Statistical significance was determined by Kruskal-Wallis with Dunn’s multiple comparisons post-test. (B) Relative mRNA level of *NSD2 and NSD3* genes was measured by RT-qPCR in head and cancer cell lines. *NSD2* and *NSD3* relative level were normalized to *18S* as a control gene. (С) Western blot of NSD2 and NSD3 protein levels in a panel of human HNSCC cell lines. (D) Quantification of Western blot images in (C). (E) mRNA level of *NSD2* and *NSD3* genes was measured by RT-qPCR after NSD1 shRNA knockdown cells at 72h after knockdown induction. *NSD2* and *NSD3* relative level were normalized to *18S* as a control gene. Statistical significance determined by ANOVA with Dunnett multiple comparison post-test. (F) Western blot of NSD2 and NSD3 protein levels upon induction of pLKO control or NSD1 shRNA knockdown at 72h with doxycycline (G) Quantification of Western blot images in (F). Statistical significance was determined by ANOVA with Dunnett multiple comparison post-test. Experiments were performed in at least three independent biological repeats. The error bars are presented as mean ± SEM. ns – not significant, *p<0.05, **p<0.01, ***p<0.001, and ****p<0.0001.

**Supplementary Figure S2.**
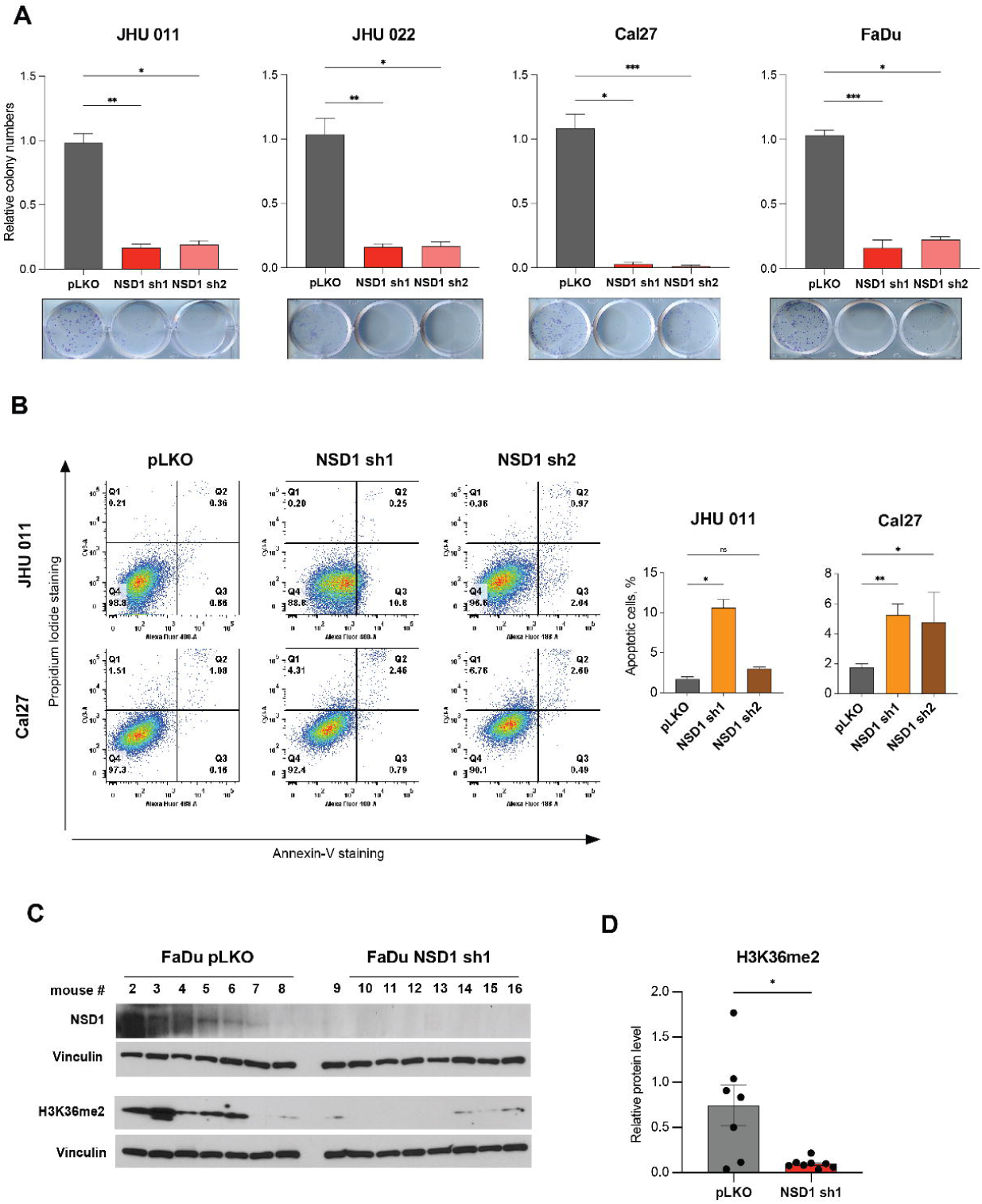
(A) Representative images of colony formation assay and quantification of the relative colony numbers in a panel of human HNSCC cell lines. Statistical significance was determined by Kruskal-Wallis with Dunn’s multiple comparisons post-test. (B) Representative flow cytometry images (left) and average calculations (right) of the Annexin V/Propidium iodide (PI) staining as measured by flow cytometry at 144h after NSD1 knockdown induction in JHU 011 and Cal27 cell lines. Statistical significance was determined by Kruskal-Wallis with Dunn’s multiple comparisons post-test. (C) Western blot of NSD1 and H3K36me^2^ levels in mice tumors. Mouse number #1 from the control group died on day 23, and the tumor of this mouse was not validated by western blot. (D) Quantification of Western blot images in (C). Statistical significance was determined by Mann-Whitney test. Experiments were performed in at least three independent biological repeats. The error bars are presented as mean ± SEM. ns – not significant, *p<0.05, **p<0.01, ***p<0.001, and ****p<0.0001.

**Supplementary Figure S3.**
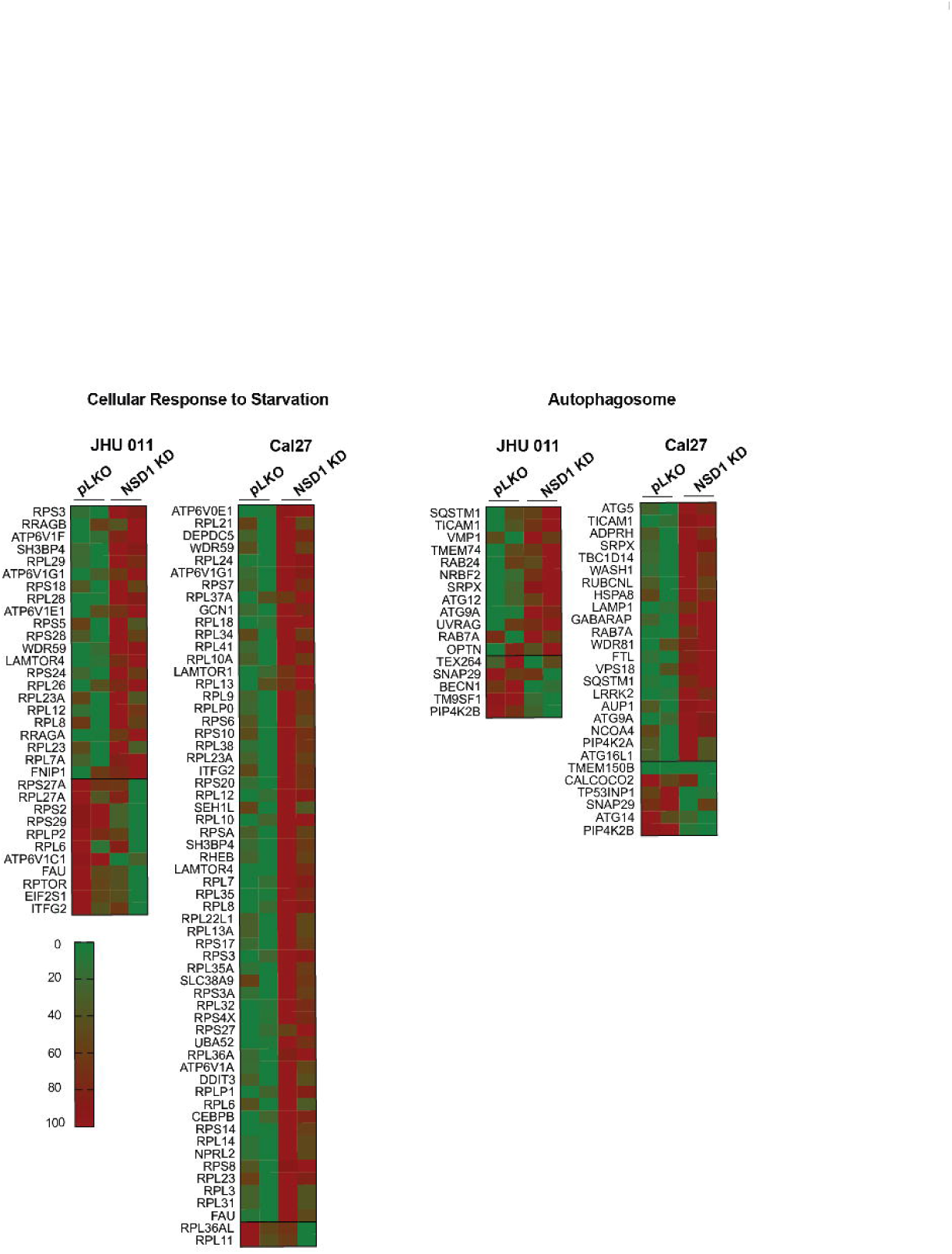
Heat maps of genes from for gene pathways from (Figure 2B).

**Supplementary Figure S4.**
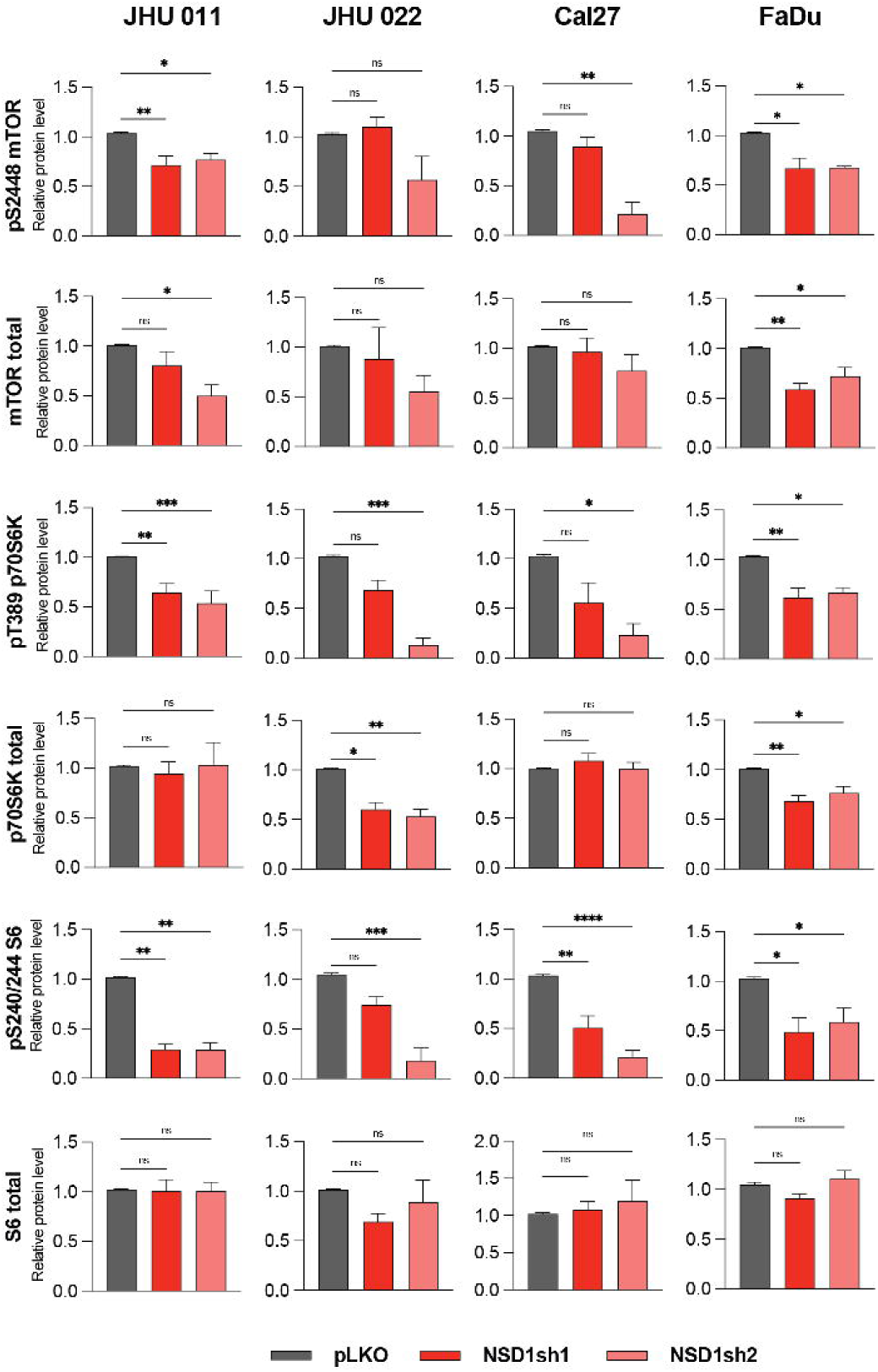
Quantification of Western blot images in Figure 3C. Statistical significance was determined by Kruskal-Wallis with Dunn’s multiple comparisons post-test. Experiments were performed in at least three independent biological repeats. The error bars are presented as mean ± SEM. ns – not significant, *p<0.05, **p<0.01, ***p<0.001, and ****p<0.0001.

**Supplementary Figure S5.**
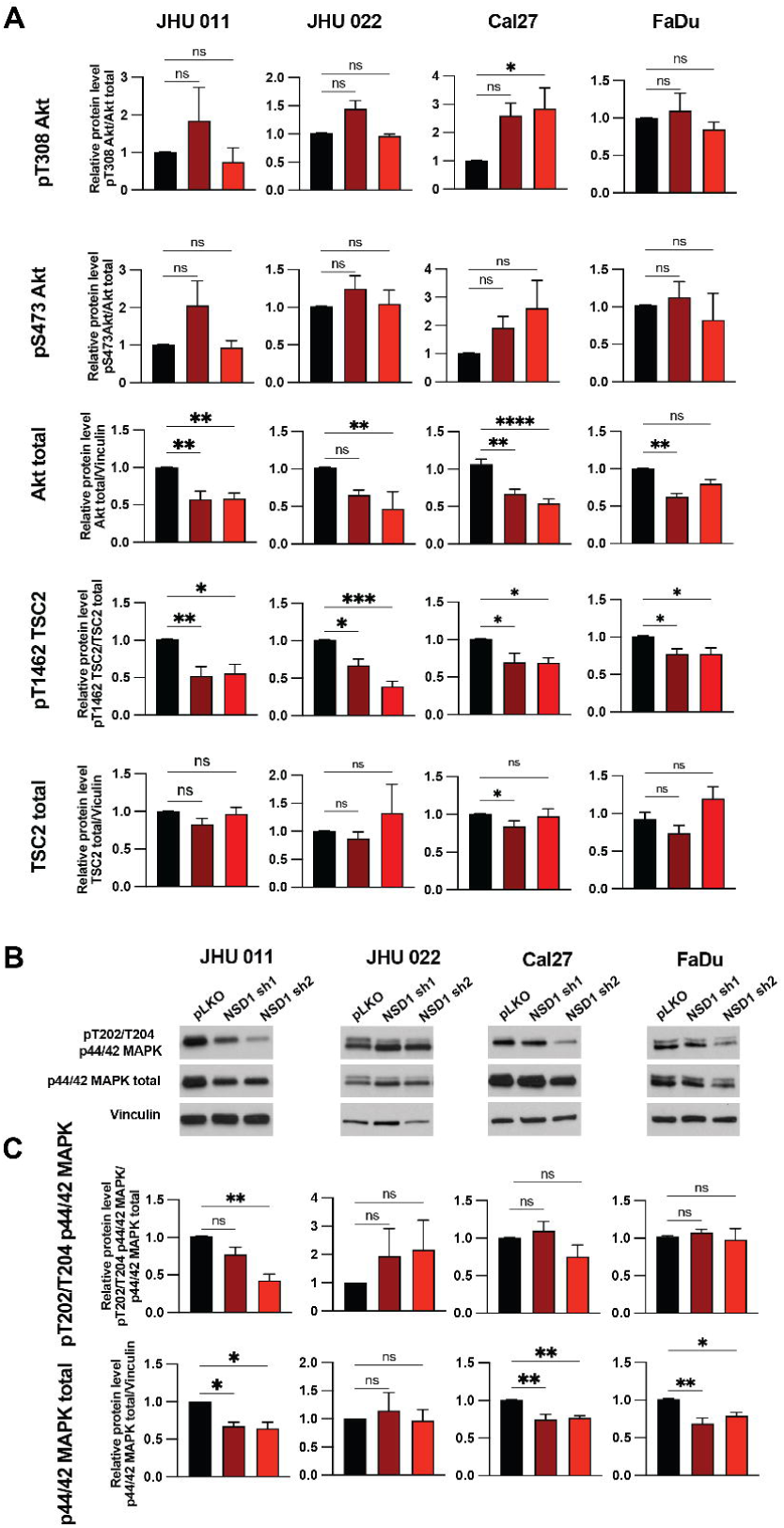
(A) Quantification of Western blot images in Figure 3D. Statistical significance was determined by Kruskal-Wallis with Dunn’s multiple comparisons post-test. (B) Western blot of pT202/204 MAPK and p44/p42 MAPK total protein levels in NSD1 shRNA knockdown cells at 72h after knockdown induction. (C) Quantification of western blot images in (B). Statistical significance was determined by Kruskal-Wallis with Dunn’s multiple comparisons post-test. Experiments were performed in at least three independent biological repeats. The error bars are presented as mean ± SEM. ns – not significant, *p<0.05, **p<0.01, ***p<0.001, and ****p<0.0001.

