## Supplementary tables for "NSD1 supports cell growth and regulates autophagy in HPV-negative head and neck squamous cell carcinoma"

**Supplementary Table 1. Cell lines characterization**

| cell line | histology type | mutations | HPV status |
|---|---|---|---|
| FaDu | Hypopharynx squamous cell carcinoma | *TP53/CDKN2A* | negative (-) |
| Cal27 | Tongue squamous cell carcinoma | *TP53/CDKN2A* | negative (-) |
| SCC61 | Tongue squamous cell carcinoma | *TP53* | negative (-) |
| SCC4 | Tongue squamous cell carcinoma | *TP53/CDKN2A/NSD1 (S752Lfs*16)* | negative (-) |
| JHU 022 | Laryngeal squamous cell carcinoma | *TP53* | negative (-) |
| JHU 011 | Laryngeal squamous cell carcinoma | *TP53* | negative (-) |

**Supplementary Table 2. shRNA oligos**

| | |
|---|---|
| NSD1 shRNA oligo 1 fw | CCGG CCGAGACGTCTCAGGTTAATCCTCGAGGATTAACCTGAGACGTCTCGGTTTTT |
| NSD1 shRNA oligo 1 rev | AATT AAAAACCGAGACGTCTCAGGTTAATCCTCGAGGATTAACCTGAGACGTCTCGG |
| NSD1 shRNA oligo 2 fw | CCGG AGGAGTGGATGGGACATATAACTCGAGTTATATGTCCCATCCACTCCTTTTTT |
| NSD1 shRNA oligo 2 rev | AATT AAAAAAGGAGTGGATGGGACATATAACTCGAGTTATATGTCCCATCCACTCCT |

**Supplementary Table 3. siRNA sequences**

| Gene symbol | Sequence | Supplier | Catalog number |
|---|---|---|---|
| NSD1 (si1) | 5`-CUCAAGAACAUGAUAUCACUAAUTT-3` | Integrated DNA Technologies | #402447171 |
| | 3`-ACGAGUUCUUGUACUAUAGUGAUUAAA-5` | | (hs.Ri.NSD1.13.1) |
| NSD1 (si2) | 5`-GUUAAAAUCAUGAAAGCAGUCACTA -3` | Integrated DNA Technologies | #402447174 |
| | 3`-ACCAAUUUUAGUACUUUCGUCAGUGAU-5` | | (hs.Ri.NSD1.13.2) |

**Supplementary Table 4. PCR primer sequences**

| Gene symbol | Primer sequence | | Supplier | RefSeqNumber |
|---|---|---|---|---|
| *NSD1* | Primer 1 | 5'-AGCTCGTCTCCTGCAAGA-3' | Integrated DNA Technologies | NM_ 172349 |
| | Primer 2 | 5'-CAGATGTCACACTGATGCCA-3' | | |
| *WHSCl (NSD2)* | Primer 1 | 5'-TCGGAAGAGAGACACAATCAC-3' | Integrated DNA Technologies | NM_ 133335 |
| | Primer 2 | 5'-GTGGTTTACATGCATCAGACAG-3' | | |
| *WHSC1L1 (NSD3)* | Primer 1 | 5'-GTATCATCTCCTGAAGCAACATC-3 | Integrated DNA Technologies | NM_023034 |
| | Primer 2 | 5'-GAACTGTTTCAACCTGCTCCT-3' | | |
| *ULK1* | Primer 1 | 5'-CTACCTGGTTATGGAGTACTGC-3' | Integrated DNA Technologies | NM_003565 |
| | Primer 2 | 5'-GGAAGAGCCTGATGGTGTC-3' | | |
| *AKT* | Primer 1 | 5'- CTCCCCTCAACAACTTCTCTG-3' | Integrated DNA Technologies | NM_005163 |
| | Primer 2 | 5'- GCGTTCGATGACAGTGGT-3' | | |
| *RNA18S5* | Primer 1 | 5'-GAGACTCTGGCATGCTAACTAG-3' | Integrated DNA Technologies | NR_003286 |
| | Primer 2 | 5'-GGACATCTAAGGGCATCACAG-3' | | |

**Supplementary Table 5. Antibodies and dilutions**

| Antibody name | Assay (IHC, Western blot, IHC) | Supplier, catalog number | Dilution |
|---|---|---|---|
| **Primary antibodies** | | | |
| NSD1 | Western blot | NeuroMab, 75-280 | 1:750 |
| NSD1 | IHC | Invitrogen, #PA5-84938 | 1:500 |
| WHSC1/NSD2 | Western blot | Abcam, ab75359 | 1:1000 |
| WHSC1L1 (NSD3) | Western blot | Cell Signaling Technology, #92056 | 1:1000 |
| Di-Methyl-Histone H3 (Lys36) | Western blot/IHC | Cell Signaling Technology, #2901 | 1:1000/1:200 |
| Histone H3 | Western blot | Cell Signaling Technology, #14269 | 1:1000 |
| Phospho-mTOR (Ser2448) | Western blot | Cell Signaling Technology, #2971 | 1:1000 |
| mTOR | Western blot | Cell Signaling Technology, #4517 | 1:500 |
| Phospho-p70 S6 Kinase (Thr389) | Western blot | Cell Signaling Technology, #9205 | 1:1000 |
| p70 S6 Kinase | Western blot | Cell Signaling Technology, #34475 | 1:1000 |
| Phospho-S6 Ribosomal Protein (Ser240/244) | Western blot | Cell Signaling Technology, #2215 | 1:1000 |
| S6 Ribosomal Protein | Western blot | Cell Signaling Technology, #2217 | 1:1000 |
| Phospho-Akt (Ser473) | Western blot | Cell Signaling Technology, #4069 | 1:2000 |
| Phospho-Akt (Thr308) | Western blot | Cell Signaling Technology, #4056 | 1:1000 |
| Akt (pan) | Western blot | Cell Signaling Technology, #2920 | 1:2000 |
| Phospho-Tuberin/TSC2 (Thr1462) | Western blot | Cell Signaling Technology, #3617 | 1:1000 |
| Tuberin/TSC2 | Western blot | Cell Signaling Technology, #4308 | 1:1000 |
| Phospho-p44/42 MAPK (Erk1/2) (Thr202/Tyr204 | Western blot | Cell Signaling Technology, #4370 | 1:2000 |
| p44/42 MAPK (Erk1/2) | Western blot | Cell Signaling Technology, #9201 | 1:1000 |
| ULK1 | Western blot | Cell Signaling Technology, #6439 | 1:700 |
| SQSTM1/p62 | Western blot | Cell Signaling Technology, #5114 | 1:1000 |
| LC3A/B | Western blot | Cell Signaling Technology, #12741 | 1:1000 |
| Phospho-AMPKα (Thr172) | Western blot | Cell Signaling Technology, #50081 | 1:1000 |
| AMPKα | Western blot | Cell Signaling Technology, #2793 | 1:700 |
| Phospho-Beclin-1 (Ser30) | Western blot | Cell Signaling Technology, #35955 | 1:1000 |
| Phospho-Beclin-1 (Ser93) | Western blot | Cell Signaling Technology, #14717 | 1:1000 |
| Beclin-1 | Western blot | Cell Signaling Technology, #4122 | 1:700 |
| Vinculin | Western blot | Cell Signaling Technology, #13901 | 1:500 |
| p62 | IHC | Novus Bio, NBP1-48320 | 1:600 |
| LC3B | IHC | Novus Bio, NB100-2220 | 1:600 |
| **Secondary antibodies** | | | |
| Anti-rabbit IgG, HRP-linked Antibody | Western blot | Cell Signaling Technology, #7074 | 1:1500 |
| Anti-mouse IgG, HRP-linked Antibody | Western blot | Cell Signaling Technology, #7076 | 1:1500 |

**Supplementary Table 6. Patient characteristics for the primary HNSCC tumors**

| **Anatomical site** | **%** |
|---|---|
| Larynx | 11 |
| Lymph node | 3 |
| Neck | 11 |
| Tongue | 67 |
| Tonsil | 8 |
| **Prognostic stage** | **%** |
| I | 0 |
| II | 11 |
| III | 36 |
| IVa | 19 |
| IVb | 33 |
| **T stage** | **%** |
| Tx | 2 |
| T1 | 8 |
| T2 | 19 |
| T3 | 44 |
| T4 | 25 |
| **Lymph nodes** | **%** |
| Negative | 36 |
| Positive | 64 |

TMAs contained specimens from 36 patients (with characteristics --noted in the table), and for 6 normal epithelial tissues.

**Table 7. PCR primer sequences for ChIP-assay**

| Gene symbol | Primer sequence | | Supplier |
|---|---|---|---|
| ULK1(-1225…-1142) | Primer 1 | 5'- CGTGTACGGTGAACAGCACT-3' | Integrated DNA Technologies |
| | Primer 2 | 5'- GGGCTCACTCACAGAAGACA-3' | |
| ULK1 (-636…-552) | Primer 1 | 5'-CCTAACCTCTAACTCAGCCATTCT-3' | Integrated DNA Technologies |
| | Primer 2 | 5'-GAGAACAGGCGTGGGAAATG-3' | |
| ULK1 (+1038…+1175) | Primer 1 | 5'-CAAAATCCTGAAGGTGAGCCAG-3 | Integrated DNA Technologies |
| | Primer 2 | 5'-GTCGTACAGGGCCACGATG-3' | |
| ULK1 (+15 319…+15 418) | Primer 1 | 5'-GCCCTACTGCAACGCAAC-3' | Integrated DNA Technologies |
| | Primer 2 | 5'-ATCAGCAAATGCAAGGAAGGAG-3' | |
| ULK1 (+28 439…+ 28 540) | Primer 1 | 5'-GTGTGATTTCCTGCCCTTTGC-3' | Integrated DNA Technologies |
| | Primer 2 | 5'-CAAGCCAACGAAGACAAGTGG-3' | |
